# Delayed recovery of seed-dispersal interactions after deforestation

**DOI:** 10.1101/2025.03.11.641853

**Authors:** A. R. Landim, J. Albrecht, J. Brito, S. Burneo, S. Erazo, B. A. Tinoco, M. Tschapka, E. L. Neuschulz, M. Schleuning

## Abstract

Interactions between plants and animals are crucial for natural forest recovery because most plants rely on seed dispersal by animals to recolonize degraded habitats. Despite this importance, the recovery time of seed-dispersal interactions after deforestation is unknown. We compared the recovery time of the functional diversity of seed-dispersal interactions, frugivorous animals and fleshy-fruited plants by recording trait and interaction data along a chronosequence of tropical forest recovery. Using a Bayesian model, we estimated that plant functional diversity was high shortly after deforestation, but animal functional diversity recovered only after 39 years, and seed-dispersal interactions after 18 years. We demonstrate that seed-dispersal interactions need about two decades to functionally recover which provides a new benchmark for the timing of restoration projects in tropical forests.

## Main Text

Deforestation and habitat degradation are decreasing the area of old-growth tropical forests (*1, 2*). Nowadays, old-growth forests with no signs of human disturbance represent only about one third of standing forests, while the other two thirds are composed of degraded or recovering secondary forests (*2*). Restoration is, thus, essential to maintain and recover tropical forest biodiversity. Traditionally, species diversity has been used to evaluate restoration success (*3*). However, the recovery of ecological processes is essential to restore ecosystem functioning in tropical forests (*4*). To assess the speed of recovery in secondary forests, it is, thus, crucial to move beyond measures of species diversity and to quantify how quickly the functional diversity of species and interactions can recover after deforestation.

Ecological processes stemming from plant-animal interactions, such as seed dispersal, are vital for plant regeneration (*5*). In tropical forests, nearly 90% of plants rely on animals for seed dispersal (*6*), making animal-mediated seed dispersal essential for plants to colonize new areas and restore degraded habitats (*7, 8*). Various frugivorous animal species, particularly birds and mammals, disperse seeds and play different roles in seed dispersal. While seed-dispersing birds are the most species-rich group of frugivores (*9*), scatter-hoarding mammals are crucial for dispersing large fruits that lay on the forest floor (*10*). The different roles of these animal seed dispersers are mediated by their functional traits and that of their partners, such as animal body mass and plant crop mass, that influence which plant and animal species interact (*11, 12*). Studying different types of animal seed dispersers and their functional traits is, thus, needed for a comprehensive understanding of animal-mediated seed dispersal in the recovery of tropical forests (*13*).

Natural forest recovery is a cost-effective restoration strategy driven by seed-dispersal interactions between plants and animals (*14, 15*). Pioneer trees can rapidly recolonize degraded habitats (*16*), enhancing the functional diversity of the plant community already within the first years of forest recovery (*17*). These pioneer plants can attract frugivores that bring seeds from other sources, such as late-successional plant species from forest remnants, promoting the regeneration process (*18*). However, seed dispersal is often constrained in degraded habitats due to a low diversity of animal seed dispersers (*19*), potentially slowing down the recovery process. Habitat degradation can further limit the availability of other resources, such as roosting sites, and hinder the movement of animal species (*20*). These factors may foster the presence of small-bodied over large-bodied animal species and reduce the diversity of animal functional traits (*21, 22*). Despite the critical role of seed dispersal in natural forest recovery, the recovery times of seed-dispersal interactions remain largely unexplored. While a previous study has investigated how seed-dispersal syndromes in a tropical plant community change over time (*23*), the time needed for the recovery of seed-dispersal interactions is unknown.

In this study, we estimate the recovery time of seed-dispersal interactions, as well as the recovery times of frugivorous animals and fleshy-fruited plants, based on an unprecedented empirical data set. Our study was conducted along a chronosequence of forest recovery in the Ecuadorian Chocó (*24*), a tropical biodiversity hotspot currently threatened by high rates of deforestation due to logging, mining, and agricultural expansion (*25*). We recorded plant-seed disperser interactions in 62 plots ranging from active pastures and cocoa plantations to recovering forests and undisturbed old-growth forests. Using a space-for-time substitution, we were able to study changes over time by comparing plots at different stages of forest recovery. To assess the recovery time of seed-dispersal interactions, we focused on three trait pairs of plants and animals related to morphological matching, energy provisioning and requirements, and foraging stratum and mobility, respectively (*26*). We hypothesized that the recovery time of the functional diversity of interactions and animals would lag behind that of plants, as the presence of plants with fruits is required for the arrival of frugivorous animals and for the establishment of seed-dispersal interactions. This potential time lag has been suggested in previous reviews (*28, 29*), but the actual time needed for the recovery of seed-dispersal interactions has never been quantified empirically.

### Functional diversity of seed-dispersal interactions

To estimate changes in the functional diversity of plants, animals and their interactions with forest recovery time, we recorded seed-dispersal interactions in three habitat types along the chronosequence: early recovery (active land use to 12 years of natural forest recovery, 19 plots), late recovery (15-38 years of natural forest recovery, 16 plots), and old-growth forests (undisturbed by human activity, 17 plots; for a detailed description of the plots see *24*). An analysis of sampling completeness showed that almost all common interactions were recorded in all habitats (fig. S1). Based on species-level trait measurements, we built functional trait spaces for fleshy-fruited plant species bearing ripe fruits, for the frugivorous animal species observed feeding on fruits, and for the actual seed-dispersal interactions (see Supplementary Methods for details). In these functional trait spaces, species and interactions were positioned according to their trait similarities, accounting for all three trait pairs (fruit diameter and gape width, crop mass and body mass, and plant height and hand-wing index, *11, 27*, fig. S2). The contribution of each interaction or species to the functional diversity of its habitat was measured by its originality, defined as the distance of an interaction or a species to its habitat’s centroid, which represents the mean position of all interactions or species observed within each of the three habitat types (Fig. 1A). Functional diversity of each plot was then calculated as functional dispersion (*30*), which is the mean originality of all interactions or species in that plot. Plots were treated as replicates to estimate functional diversity within each habitat type (Fig. 1B). Based on these data, we estimated the recovery time of functional diversity of seed-dispersal interactions, frugivorous animals and fleshy-fruited plants (Supplementary Methods). To estimate recovery time, we applied a Bayesian approach, which accounted for the variance in functional diversity both in the active agricultural land (prior to recovery) and in the reference old-growth forests (table S1). We used this model to estimate recovery rates for interactions, animals and plants and estimated the time required to reach 90% of the functional diversity observed in old-growth forests (as in *29*).

**Fig. 1.**
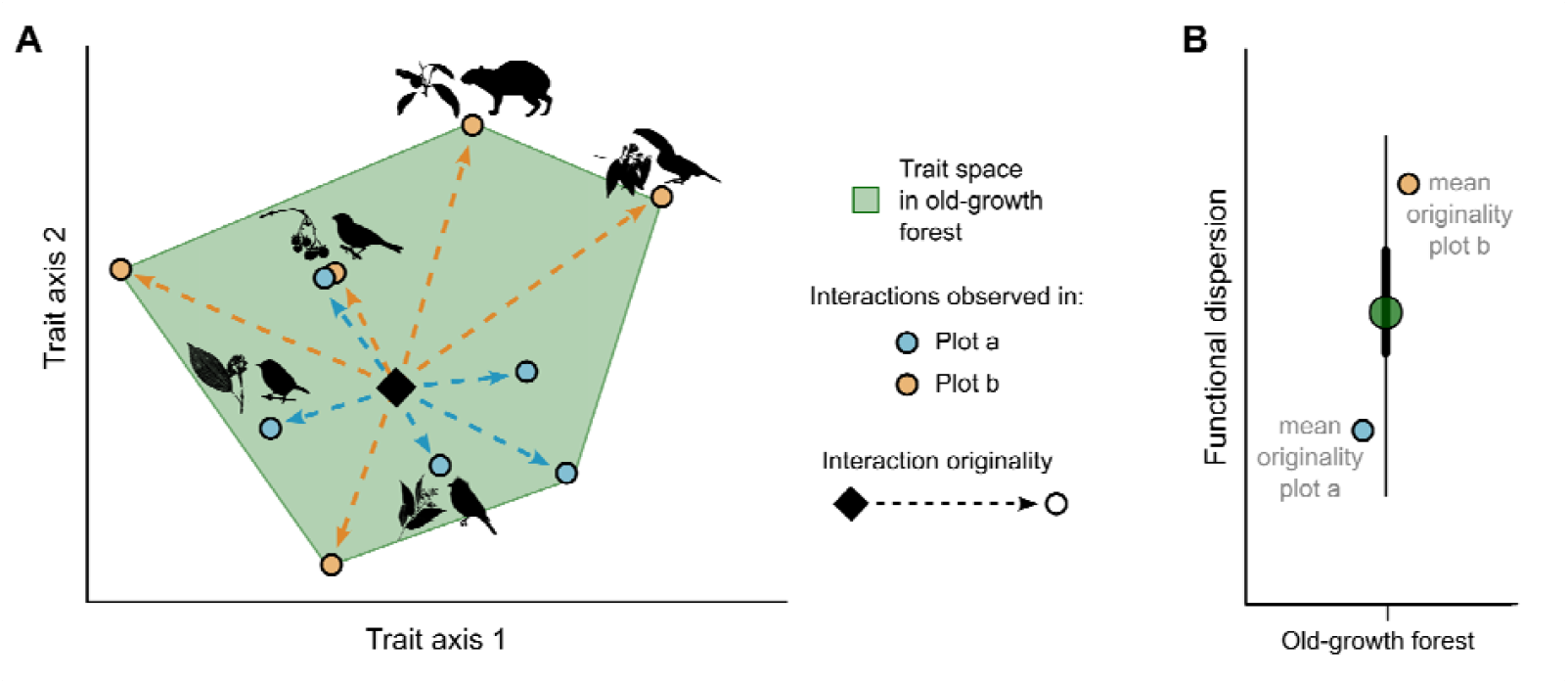
Functional diversity of seed-dispersal interactions. **(A)** Originality of interactions. Trait values are assigned to each interaction based on the traits of the interacting plant and animal species. Similar interactions cluster together, e.g., passerine birds with small-fruited plants versus toucans or agoutis interacting with large-fruited plants. The habitat centroid (black diamond) represents the mean position of all interactions in a habitat type (e.g., old-growth forest). The distance from an interaction to the centroid determines its originality. **(B)** Functional dispersion in a habitat type (e.g., median, 50 and 95% confidence intervals) is given by the mean originality of interactions in each of the plots. In the example, plot *b* has more original interactions and, thus, a higher functional diversity than plot *a*. Originality and functional dispersion were analogously estimated for plants and animals, respectively.

### Recovery time of functional diversity

Functional diversity of seed-dispersal interactions and frugivorous animals was highest in old-growth forests, while the functional diversity of fleshy-fruited plants remained relatively constant across the chronosequence (Fig. 2). In early recovery forests, most interactions occurred between small, passerine birds and small-fruited plants (Fig. 2A). Functional diversity increased in late-recovery forests as bird species with distinct traits (e.g., large gape width and pointed wings), such as toucans and doves, interacted with both pioneer plants, such as species from the genus *Cecropia*, and late-successional ones, such as *Ficus* and *Nectandra* (species names in table S2). Old-growth forests had the highest values of functional diversity, hosting mostly original interactions, such as those between large mammals and large-fruited plants. Analogous to the trend observed in interactions, the increase in animal functional diversity from early-to late-recovery forests was primarily driven by shifts in the trait space from small, passerine birds, such as tanagers and manakins in early recovery stages, to large birds and mammals in late-recovery and old-growth forests (Fig. 2B). In contrast, plant functional diversity remained rather constant across habitat types, largely due to the presence of large remnant trees throughout the chronosequence (Fig. 2C). Nevertheless, old-growth forests harbored the tree species with the largest fruit sizes and crop masses, such as *Carapa guianensis* (Meliaceae) and *Osteophloeum platyspermum* (Myristicaceae), which were not observed in recovering forests.

**Fig. 2.**
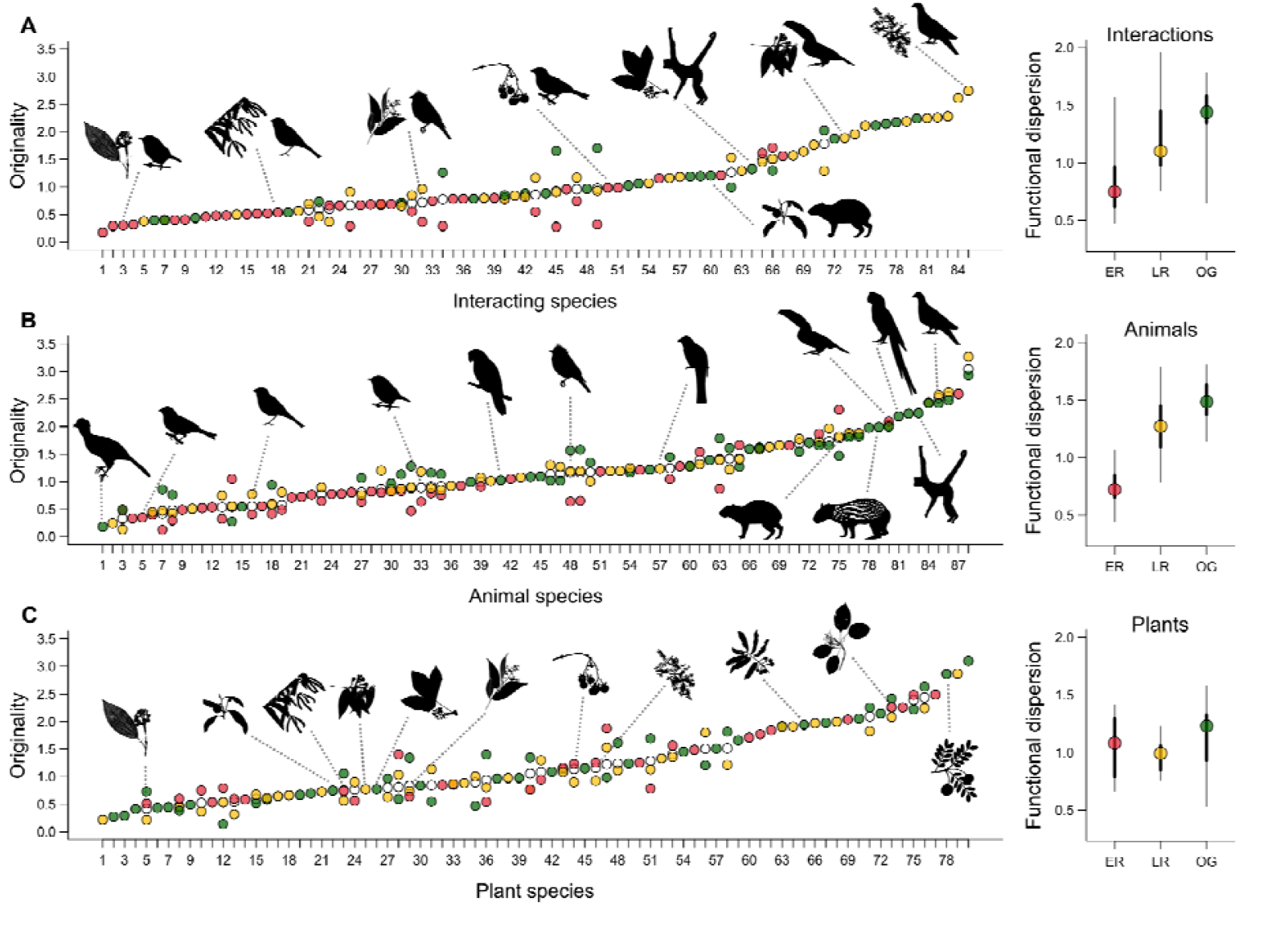
Functional diversity in recovering and old-growth forests. Originality and functional dispersion of **(A)** seed-dispersal interactions, **(B)** frugivorous animal species, and **(C)** fleshy-fruited plant species. Originality (left) represents the individual contributions of interactions or species to the functional trait space of their respective habitat types: red for early recovery (0-12 years, ER), yellow for late recovery (15-38 years, LR) and green for old-growth forest (no human disturbance, OG). White circles indicate the mean originality of interactions and species observed in more than one habitat type. In (A), only interactions observed at least six times are shown, but the entire range of originality values is covered. Functional dispersion (right) was calculated as the mean originality per plot. Dots represent the median functional dispersion, with error bars indicating the 50% (thick line) and 95% (thin line) confidence intervals across plots within each habitat type. Linear models indicated significant differences among habitat types for interactions (F_2,49_ = 14.84, p < 0.001, R^2^ = 0.38) and animals (F_2,49_ = 52.78, p < 0.001, R^2^ = 0.68), but no significant difference for plants (F_2,49_ = 2.03, p = 0.14, R^2^ = 0.07). Silhouettes of plants and animals represent important interactions and species (copyrights of animal silhouettes: phylopic.org, plant silhouettes were created by A.R.L.; see tables S2-4 in the Supplementary Material for full species names and trait values).

Seed-dispersal interactions, frugivorous animals and fleshy-fruited plants had distinct recovery trajectories following deforestation (Fig. 3). Animals took much longer than plants to reach 90% of the functional diversity observed in old-growth forests (Fig. 3D). Consequently, the functional diversity of interactions lagged behind the rapid recovery of plant functional diversity. The functional diversity of interactions required approximately 18 [1, 52] years (median [95% credible interval]) to recover. In comparison, animal functional diversity required 39 [23, 62] years, whereas plant functional diversity was already after one [1,2] year of recovery similarly high as in the old-growth forests.

**Fig. 3.**
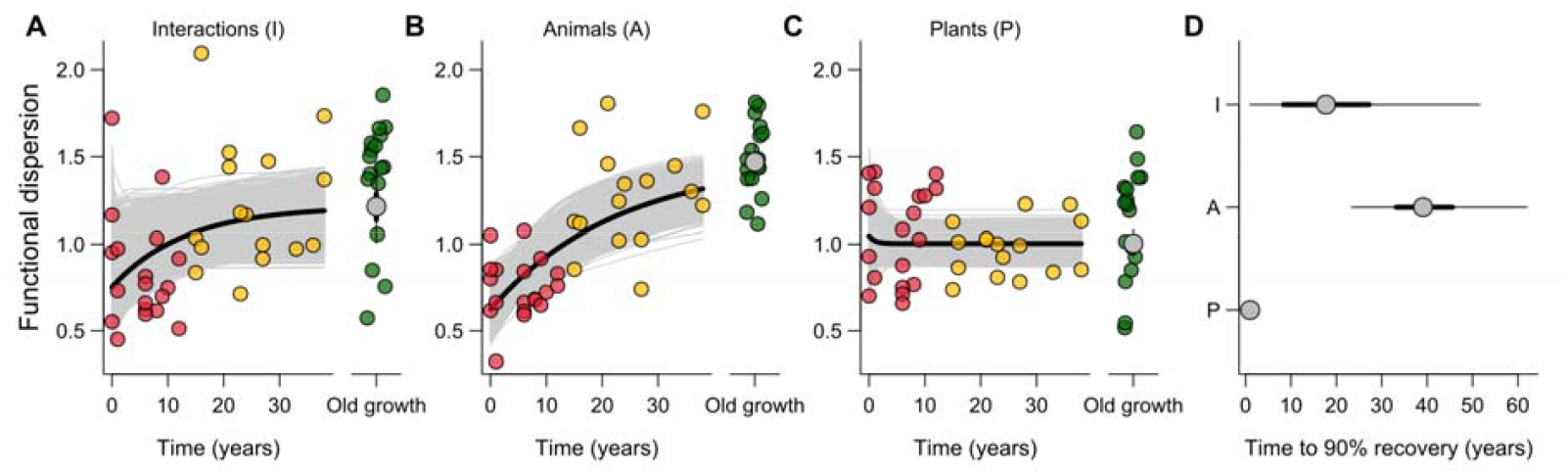
Recovery time of functional diversity of seed-dispersal interactions, animals and plants. **(A to C)** Recovery trends of the functional dispersion of (A) seed-dispersal interactions, (B) frugivorous animals and (C) fleshy-fruited plants. Red, yellow and green points represent functional dispersion in early recovery, late recovery, and old-growth forests, respectively. Gray lines indicate recovery trajectories predicted by the models; the black line is the median prediction. Functional dispersion observed in old-growth forests was used as a reference for recovery, with gray points indicating median values, and thick and thin lines representing the 50% and 95% credible intervals (CrI) from the Bayesian posterior distribution (modeled with a log-normal distribution). **(D)** Estimated recovery time to reach 90% of old-growth forest functional dispersion. Functional diversity of interactions tended to recover faster than that of animals [p(*t*_90_I < *t*_90_A) = 0.91, posterior probability] and slower than that of plants [p(*t*_90_I > *t*_90_P) = 0.88]. Animal functional diversity recovered slower than plant functional diversity [p(*t*_90_A > *t*_90_P) = 1].

Our analysis supports our hypothesis that animal functional diversity recovers slower than that of plants, thus delaying the recovery of seed-dispersal interactions. Nevertheless, the recovery of seed-dispersal interactions occurred faster than that of frugivorous animals alone. One reason for this is the early availability of fleshy-fruited plants due to the presence of remnant trees with original traits, such as large fruits or crop mass [e.g., *Coussarea latifolia* (Rubiaceae) and *Otoba novogranatensis* (Myristicaceae)]. Early recovery plots harboring these remnant trees had a high functional plant diversity and attracted functionally distinct frugivore species despite their short recovery time (Fig. 3C). By attracting frugivore species which are otherwise rare in degraded habitats (*21, 32*), remnant trees can facilitate the dispersal of a larger diversity of seeds from forest fragments into early-recovery forests. For example, rodents were the only animal dispersers of remnant trees with medium and large fruits like *Wettinia quinaria* (Arecaceae) and *Grias peruviana* (Lecythidaceae) in early-recovery forests. The presence of remnant trees is therefore key for forest recovery because they can promote the dispersal of a larger diversity of seeds from forest fragments into abandoned agricultural land and secondary forests (*33, 34*).

We found slow recovery times of the functional diversity of interactions and animals (Fig. 3A and B). The main reason for this was that the interactions involving animal species with distinct traits appeared rather late along the chronosequence of forest recovery. Specifically, most interactions involving large-gaped animals and birds with pointed wings were only observed after 20 years of forest recovery. Large-gaped animals can interact with larger fruits, thereby contributing differently to seed dispersal compared to small-gaped animals (*35*). Furthermore, animals with pointed wings have greater mobility, enhancing the chances of seed dispersal over longer distances and between old-growth forest fragments and recovering forests (*21*). Additionally, interactions involving most of the large animal species (e.g., Baudó guans and white-lipped peccaries) were almost only observed in old-growth forests. Large-bodied animals can handle different-sized fruits, disperse seeds over long distances (*36*), consume high quantities of fruit and tend to handle fruits in ways that facilitate plant regeneration, which makes them particularly effective seed dispersers (*37*). The absence of interactions involving these large-bodied animals has therefore been associated with reduced seedling diversity and the local extinction of plant species (*36, 38*). In our study, the lack of most of these large-bodied animal species in early and late recovery forests explains why seed-dispersal interactions recovered more slowly than plant functional diversity.

We found that seed-dispersal interactions involving highly mobile animal species such as parrots and spider monkeys were absent in recovering forests even though these forests were mostly in spatial proximity to the old-growth forest (mean distance between recovering and old-growth forests: 58±7 m, *24*). This suggests that animal movement was not a primary factor limiting the recovery of seed-dispersal interactions. Furthermore, although fruit availability has been identified as a key driver of frugivore diversity in degraded habitats (*34, 39, 40*), this is unlikely to be a major limiting factor in our study because fruit abundance and diversity showed no significant trends along the chronosequence of forest recovery (fig. S3). Instead, other factors may have constrained seed-dispersal interactions involving certain animal species in recovering forests. One reason could be that animal seed dispersers changed their feeding habits in the degraded and restoring habitats, e.g., by consuming agricultural resources when those are available (*41*). In our study area, pacas consumed native plants in old-growth forests, whereas they fed exclusively on cocoa fruits in recovering forests. In addition, hunting pressure may have also played a significant role, as recovering forests were more accessible to humans than old-growth forests. Large-bodied frugivores are particularly susceptible to hunting and have lower abundances in areas with higher hunting intensity (*42*). Finally, studies have shown that plant community composition and above-ground biomass can take up to 120 years to recover (*31*), suggesting that the structural complexity of tropical forests will only be fully recovered after more than a century. Such long recovery times may limit the presence of certain animal species from recovering forests, due to the lack of suitable nesting or shelter sites related to the presence of specific tree species (*28*). Some species, such as the long-wattled umbrellabird, prefer to feed in old-growth forests with closed canopy (*43*) and rely on densely forested areas used as leks for mating (*44*).

Current strategies of forest restoration often prioritize to attract animal seed dispersers in the first years after deforestation (*22, 34, 45*). According to our findings, this may be insufficient if only a subset of animal seed dispersers, and the seeds they carry, can be attracted to young sites. This particularly applies to large animals with specific habitat requirements. Restoration management should foster the return of such species by considering the specific requirements of these species related to nesting, shelter and foraging (*45*). This aligns with recent studies showing that planting diverse tree islands can accelerate the recovery of tropical forests by fostering the dispersal of a more diverse range of plant species (*46*). Such targeted restoration measures may need more time and resources, but may be critical to facilitate the return of animal species that are able to disperse late-successional plant species.

Our study was conducted under conditions that are highly favorable to natural forest recovery because most recovering sites were connected and located in close proximity to old-growth forests. In more fragmented landscapes, natural forest recovery may take considerably longer. For example, while in our study toucans started to contribute to seed-dispersal interactions within a decade after deforestation, another study reported that they were still absent after 25 years in a highly fragmented forest (*47*). Another limiting factor could be a low vegetation cover that can impede the movement of large animals to regenerating areas (*22, 48*). In such landscapes, most seed-dispersal events are performed by small birds over short distances, leading to a low connectivity and a slow recovery of tropical forests (*48*). Compared to our study, these previous studies suggest that recovery times may be substantially longer in more isolated and degraded landscapes. It is therefore likely that the estimated recovery time of about two decades may define the minimum time required for the functional recovery of seed-dispersal interactions in tropical forests.

## Conclusions

Most tropical secondary forests are composed of patches with less than 10 to 15 years of recovery age (*49*) because many of these patches are cleared again after this time period for agricultural use (*50*). Our study shows that this is insufficient time for frugivorous animals and seed-dispersal interactions to re-establish, which is a critical limit to the recovery of biodiversity and ecosystem functioning in tropical forests. Our finding calls for long-term strategies that allow secondary forests to mature and recover the ecological processes required to underpin natural forest recovery. Specifically, our findings demonstrate that management programs of secondary tropical forests would need to span at least two decades to facilitate the recovery of seed-dispersal functions. Although this requires long-term strategies in conservation management, this time scale is within a human’s lifetime, which makes it a feasible target for the restoration of tropical forests.

## Supporting information

Supplementary Methods and Results

## Acknowledgments

We thank the Fundación Jocotoco and Fundación Reserva Tesoro Escondido for logistical support and permission to conduct research in their reserves, especially Martin Schaefer and Citlalli Morelos. We would like to thank especially Jordi Ninabanda and Jefferson Tacuri for support in data collection, and Katrin Krauth, Bryan Tamayo, Julio Carvajal, Yadira Giler, Franklin Quintero, Lady Condoy, Leonardo de la Cruz, Segundo Tacuri, Vicente Vélez and Vinicio Vélez for additional support and field assistance. We thank Nico Blüthgen, María-José Endara, Constance J. Tremlett, and Edith Villa-Galaviz for project coordination and administration. We acknowledge the Ministerio del Ambiente, Agua y Transición Ecológica, Ecuador for granting collection and research permits under the Genetic Resources Access Agreement number “MAATE-DBI-CM-2021-0187.”

## Funding

Deutsche Forschungsgemeinschaft (DFG), Research Unit REASSEMBLY (FOR 5207; sub-project SCHL 1934/5-1).

Deutsche Forschungsgemeinschaft (DFG), Research Unit REASSEMBLY (FOR 5207; sub-project NE 1863/4-1).

Deutsche Forschungsgemeinschaft (DFG), Research Unit REASSEMBLY (FOR 5207; sub-project MT TS81/15-1).

## Author contributions

Conceptualization: ARL, ELN, MS

Data collection: ARL, SE, JB

Analysis: ARL, JA

Visualization: ARL

Funding acquisition: ELN, MS, MT

Project supervision: ELN, MS, MT, BT, SB

Writing – Original Draft: ARL, ELN, MS

Writing – Review & Editing: all authors

## Data and materials availability

https://github.com/annarlandim/Recovery-of-seed-dispersal-interactions

