## Supplementary Methods and Results for "Delayed recovery of seed-dispersal interactions after deforestation"

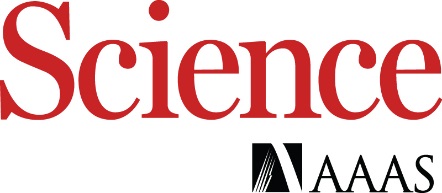

Supplementary Materials for

**Delayed recovery of seed-dispersal interactions after deforestation**

A. R. Landim^1,2^*, J. Albrecht^3,4^, J. Brito^5^, S. Burneo^6^, S. Erazo^6,7^, B. Tinoco^8^, M. Tschapka^7,9^, E. L. Neuschulz^1^† and M. Schleuning^1^†

**The PDF file includes:**

Materials and Methods

Figs. S1 to S3

Tables S1 to S4

References

Materials and Methods

All analyses were done using R version 4.3.1.

Study area

The study was conducted in the Chocó lowland forest of northern Ecuador, a biodiversity hotspot currently under threat from extensive deforestation (*25*). The region’s climate is characteristic for equatorial lowland rainforests, with an annual rainfall of approximately 5000 mm, and temperature ranging from 21 to 25°C (*24*). The study was carried out within and around the Canandé and Tesoro Escondido private reserves (0.5°N 79.2°W, 130–540 masl), covering 14,000 and 1,800 hectares, respectively. Over the last 50 years, the land has been subjected to rapid transformation, including logging of old-growth forests for timber and conversion into cacao plantations, oil palms, and pastures for livestock. Much of this land has since then been acquired by the reserves and was left to recover, forming a mosaic of degraded and recovering habitats, as well as old-growth forests.

Plot design

We recorded seed-dispersal interactions at 62 plots (50 m x 50 m each) along a forest recovery gradient including 19 plots of early recovery forests (active agricultural land to 12 years of recovery), 16 plots of late recovery forests (15-38 years of forest recovery), and 17 plots of old-growth forests. Plots were selected in the framework of the Reassembly project (DFG FOR 5207, https://www.reassembly.de) and habitat categories were set based on the structural characteristics of the forest (*24*). Plots were selected to maximize spatial independence, with distances of at least 180 m between plots of similar recovery age, minimizing biases from surrounding forest cover, elevation or land use (for specific details on the plot design please refer to Escobar et al. *24*). By using a space-for-time substitution, we were able to assess the recovery of seed-dispersal interactions over time.

Seed-dispersal interactions

To record seed-dispersal interactions across all forest strata, we used two different methods at each of the 62 plots: direct observations and camera-trap recordings. Sampling via both methods was done simultaneously and carried out from 2022 to 2023 during rainy (March-June) and dry (September-December) seasons. The sampling of plots within the three the habitat types was equally distributed across rainy and dry seasons.

For the upper forest layers (midstory and canopy), we conducted direct observations of seed-dispersal interactions of birds and mammals using binoculars. Observations were conducted for 5 hours starting at sunrise over three consecutive days, resulting in 15 hours of observations per plot. In very rare situations, we had to pause observations because of heavy rain. Although the largest trees reached over 50 meters in height, the steep slopes of the study area and animal movement within the plot facilitated species identification. The few observations that could not be assigned to species were discarded from the analyses. Birds were identified following Restall & Freile (*51*) and mammals following Tirira (*52*).

On the forest floor, we recorded interactions by deploying fruits in front of four motion-triggered camera traps placed at each corner of the plots. Fruits from all fruiting plants within a plot were deployed in front the cameras. Deployed fruits were collected within the plot and were, when necessary, supplemented with fruits of the same species available in the surrounding areas. Out of the 81 plant species recorded in this study, 54 species could be placed in front of the camera traps. Cameras ran continuously for six days and captured 30-second videos with 1-second intervals between the recordings. Bird species were identified following Restall & Freile (*51*) and mammals following Tirira (*52*).

In both methods, we recorded all occasions where animals interacted with fleshy fruits, either by swallowing (67% of interactions observed via direct observations, 26% via camera traps), pecking or biting (28% of direct observations, 54% via camera traps) or transporting entire fruits or seeds (5% of direct observations, 20% via camera traps). As most of the observed pair-wise interactions occurred in different of these categories (e.g., swallowing and pecking), we included all types of interactions to build a binary interaction matrix between plant and animal species for each plot. This matrix indicated whether an interaction occurred, regardless of its type or frequency. We removed interactions involving cultivated species, such as cacao, yucca, and jackfruit, to focus specifically on the recovery of seed-dispersal interactions that support natural forest recovery. Using information on the frequency of the observed interactions, we conducted a sampling completeness analysis using the package *iNext.link* (version 1.0.1) (*53*). We did this separately for seed-dispersal interactions recorded by direct observations and camera traps because the interaction frequencies were not directly comparable between the two methods. This analysis showed that almost all common interactions were captured by both methods (fig. S1), with approximately 95% of all interactions recorded via observations in all habitat types, and about 70% via camera traps on the forest floor.

Plant and animal traits

We collected functional trait data of the fruiting plant and frugivorous animal species observed interacting, focusing on traits relevant for trait matching in seed-dispersal interactions: gape and fruit width, avian hand-wing index and plant height, and body mass and crop mass (*11, 54*).

Bird traits (*55*) and mammal body mass (*56, 57*) were obtained from literature. Gape width of mammals was measured directly from museum collections in Quito (Museo de Zoologia, PUCE and Museo de Historia Natural, EPN). It was measured as the greatest distance between the angles of the mandible, with measurements taken from four specimens per species (except for *Cebus capucinus*, for which only one specimen was available). A list of all animal species and their traits is available in table S3.

Plant traits were measured directly in the field. Plant height and number of fruits were measured for all fruiting individuals in each plot and fruit width and mass were measured for 10 fruits per individual, sampling up to 10 individuals per species when possible. When the total number of fruiting individuals in the plots was too low, measurements were complemented with data from individuals in the surrounding area. Species-level means of all traits were calculated for each plant species. Crop mass was calculated by multiplying the mean number of fruits per species by mean fruit weight. A list of all plant species and their traits is available in table S2.

Prior to the analysis, we checked for potential collinearity between traits. We therefore normalized gape width by body mass and fruit width by crop mass, using residuals from log-log regressions (following *58*). This allowed us to unmask potential collinearities between traits while controlling for the effects of animal and fruit mass.

Analysis of functional diversity

We constructed functional trait spaces for seed-dispersal interactions, the interacting animal species and the fruiting plant species (fig. S1). These functional trait spaces were based on trait similarities, where species or interactions were positioned according to their trait values while considering all three trait pairs (gape width-fruit width, hand-wing index-plant height and body mass-crop mass). Trait values were assigned to each interaction based on the traits of the interacting plant and animal species. In that sense, each plant and animal species were characterized by three traits, while plant-animal interactions were defined by six traits.

We conducted Principal Components Analyses (PCA) to define the functional trait space of interactions, animals and plants. Given that our dataset included missing values (e.g., hand-wing index for primates and terrestrial mammals), we used pairwise deletion to compute the correlation matrix required for the PCA, employing the ‘*principal’* function from the *psych* package (version 2.4.1, *59*). Before performing the PCA, we determined the number of principal components necessary to describe the correlation matrix of the traits adequately. To do so, we calculated the correlation matrix between traits and applied a permutation approach to assess which components had eigenvalues greater than expected by chance (*58*). Specifically, we permuted the trait values while maintaining their mean and variance but breaking their correlations. By repeating this process 1,000 times, we generated a null distribution of eigenvalues for comparison against the observed eigenvalues. We selected only those components with eigenvalues larger than expected from the null distribution for the PCA. In all cases (plants, animals, interactions), two components were sufficient.

With the selected principal components, we constructed trait spaces that projected interactions or species based on their trait values onto two axes (*11, 27*). For each plot, we calculated the originality of individual interactions or species using the Euclidean distance between each interaction or species relative to the habitat centroid. The habitat centroid represents the mean trait value of all interactions or species observed in the respective habitat type. Functional diversity of each plot was then calculated as functional dispersion (*30*), measured by the mean originality of all interactions or species observed in that plot. Plot-level estimates were used as replicates for each habitat type (mean and 95% confidence intervals) and were used to compare functional diversity among early recovery, late recovery and old-growth forests.

Analysis of recovery time

To estimate the recovery time of functional diversity, we used a Bayesian hierarchical model using the *rjags* package (version 4-15) (*60*). Based on Poorter et al. (*31*), this model aimed to quantify the recovery times of the functional diversity of seed-dispersal interactions, animals, and plants to reach levels comparable to those observed in old-growth forests. The model predicted functional diversity of recovering forests as a function of recovery time, starting from active land use (i.e., recovery time equals to 0) and increasing towards an asymptote represented by the functional diversity in old-growth forests: $A_{t} = A_{0} + (A_{OGF} - A_{0}) \times(1-e^{(-\lambda*t)})$. Where $A_{t}$ represents the functional diversity of interactions, animals or plants at the time t, $A_{0}$ functional diversity a time 0, $A_{OGF}$ functional diversity observed in old-growth forests and λ the intrinsic recovery rate. In that sense, functional diversity in recovering forests was modelled as changing (either increasing or decreasing) towards old-growth forests over time, with the rate of change driven by the intrinsic recovery rate (λ).

Since our response variables (functional diversity of seed-dispersal interactions, animals, and plants) are positive by definition, we worked with underlying lognormal distributions. Posterior distributions for model parameters were obtained using Markov Chain Monte Carlo (MCMC) sampling. We ran five parallel chains for the models, with each chain running for 200,000 iterations and with thinning intervals of 200 iterations to reduce autocorrelation in the posterior samples, which resulted in 1,000 iterations per chain. Convergence of the chains was assessed through standard diagnostic measures, including Gelman-Rubin statistics (Potential scale reduction factor, PSRF) and effective sample size (N_eff_; table S1).

Using the posterior distributions of model parameters (Θ_0_, Θ_OGF_ and λ), we estimated the recovery times of the functional diversity of interactions, animals and plants as the time at which functional diversity reached 90% of the values observed in old-growth forests ($A_{OGF}$, *31*). We calculated the recovery time as the first point (t_90_) at which the modelled functional diversity exceeded 90% of the asymptotic value for increasing trends or dropped below 110% for decreasing trends, which ensured that recovery time reflected changes towards old-growth forest values. For each group, the 95% and 50 % credible intervals were calculated by summarizing the posterior distributions of t_90_. Lastly, we conducted pairwise comparisons on these values, quantifying the likelihood that the recovery time of one group exceeded another’s (e.g., animal versus plant functional diversity).

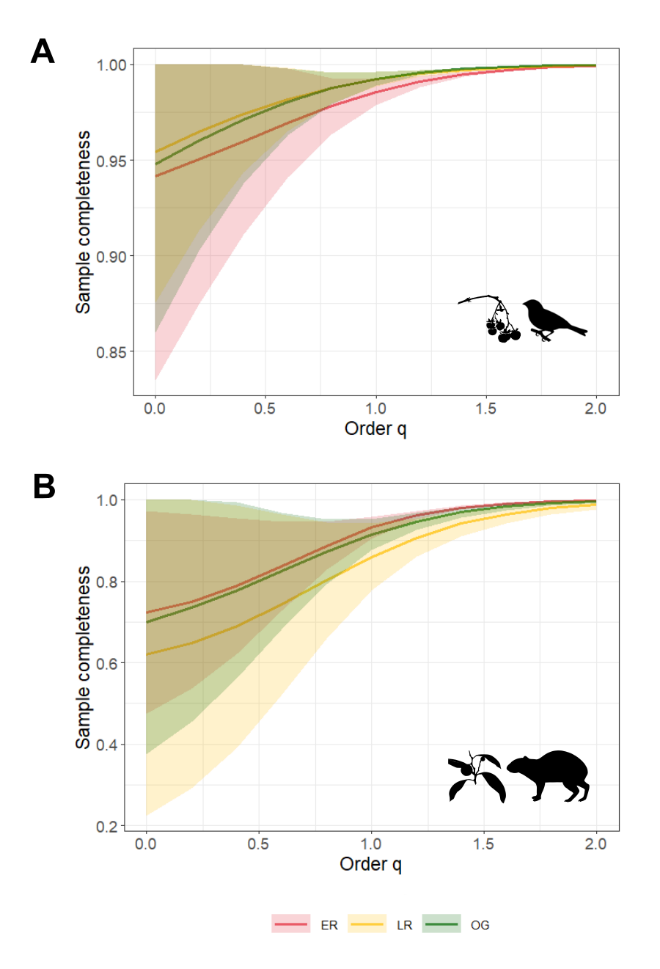

Fig. S1. Sampling completeness of seed-dispersal interactions. Shown are trends for seed-dispersal interactions based on the frequency of observed interactions for (A) direct observations and (B) camera traps. The order *q* shown on the x-axis adjusts the sensitivity of the diversity measurement to the frequency of interactions (*53*). When q=0, interaction richness is calculated (i.e., interaction frequency is not considered). When q=1, Shannon diversity is calculated, giving equal weight to rare and common interactions, and when q=2, Simpson diversity is calculated, placing more weight on frequent interactions. Shades represent 95% confidence intervals obtained via 100 bootstrap replications. Sampling completeness of seed-dispersal interactions were estimated using the *Completeness.link* and *ggCompleteness.link* functions from the *iNEXT.link* package (version 1.0.1, *53*). Color codes are the same as in the other figures: red for early recovery, yellow for late recovery and green for old-growth forests.

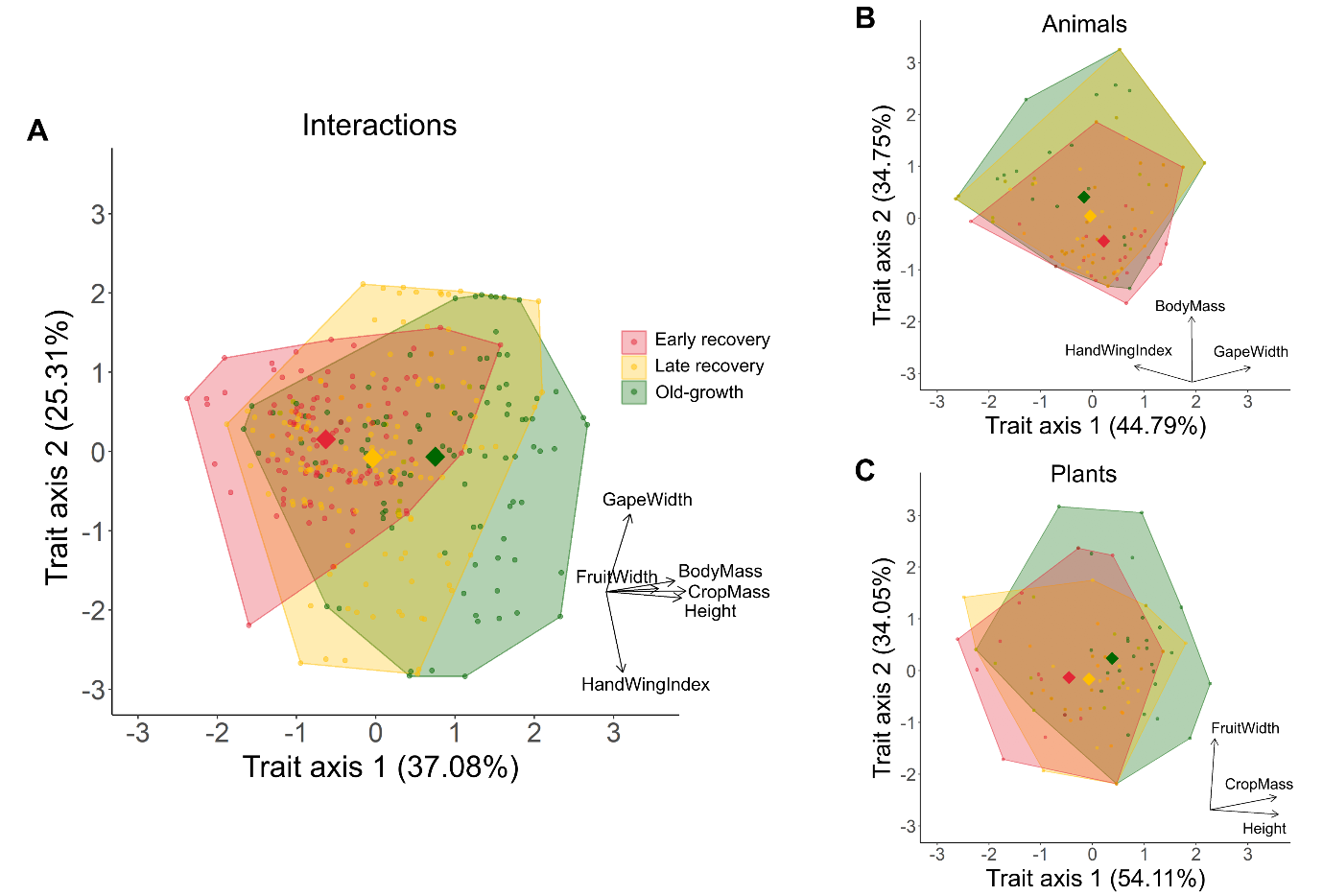
Fig. S2. Functional trait spaces of seed-dispersal interactions, animals, and fruiting plants. (A) Interactions and (B and C) species are plotted according to their trait values, with each point representing an interaction or species in trait space. Similar interactions or species cluster together, with habitat centroids (colored diamonds) representing the mean trait position for each habitat type. For all groups, two principal components were sufficient to accurately represent the trait variation. Arrows in the bottom right display the directions and loadings of the traits. For example, in the animals’ functional trait space (B), animal species with greater values of hand-wing index (i.e., more pointed wings), body mass and gape width are depicted in the left, top and right of the plot, respectively.

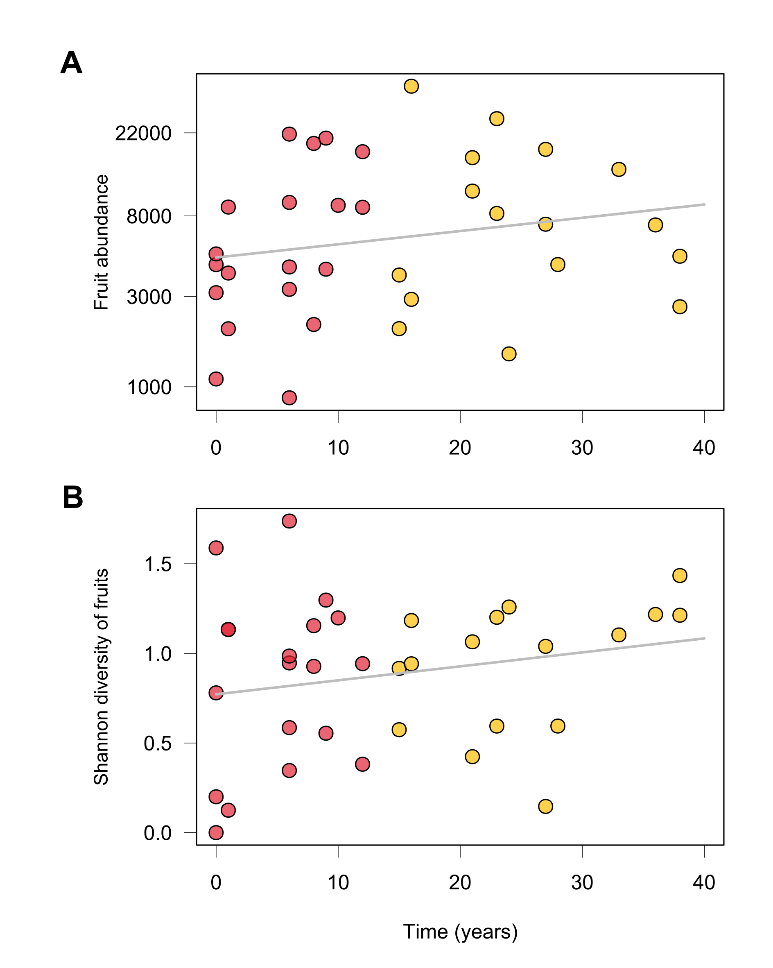

**Fig. S3. Abundance and diversity of fruits along the chronosequence of forest recovery. (A)** Fruit abundance (log-scale) and **(B)** Shannon diversity of fruits plotted against forest recovery time. Points are colored according to their habitat type: red for early recovery (0–12 years) and yellow for late recovery (15–38 years). Lines illustrate trends in fruit abundance and diversity along the chronosequence. Trend lines were fitted using generalized additive models using the *mgcv* package (version 1.8-42, *61*), which revealed no significant trend for fruit abundance (p = 0.25, adjusted R^2^ = 0.01) or fruit diversity (p = 0.22, adjusted R^2^ = 0.02).

Table S1. Diagnostics of the Bayesian hierarchical model to estimate recovery times of seed-dispersal interactions, animal, and plant functional diversity. Parameters include functional diversity values during active land use (Θ_0_), functional diversity values in old-growth forests (Θ_OG_), recovery rates (λ), and precision terms for observations in recovering (Τ_REC_) and old-growth (Τ_OG_) forests. The potential scale reduction factor (PSRF) and effective sample size (N_eff_) values indicate convergence and sampling efficiency for each parameter. A PSRF close to 1 indicates similar variability between and within Markov chains, indicating chains converged to the same distribution. N_eff_ quantifies the number of independent samples per chain.

|  | **Effect size** | | | | |  |  |
| --- | --- | --- | --- | --- | --- | --- | --- |
| **Parameter** | **Median** | **2.5%** | **25%** | **75%** | **97.5%** | **N_eff_** | **PSRF** |
| Θ_0_ interactions | 0.75 | 0.58 | 0.68 | 0.82 | 1.09 | 4098.99 | 1 |
| Θ_OG_ interactions | 1.22 | 1.01 | 1.14 | 1.29 | 1.43 | 4813.38 | 1 |
| λ interactions | 0.08 | 0.03 | 0.05 | 0.14 | 2.06 | 3128.50 | 1 |
| Τ_OG_ interactions | 0.11 | 0.06 | 0.09 | 0.15 | 0.25 | 5399.56 | 1 |
| Τ_REC_ interactions | 0.13 | 0.08 | 0.11 | 0.16 | 0.23 | 5212.84 | 1 |
| Θ_0_ animals | 0.62 | 0.51 | 0.58 | 0.67 | 0.75 | 5690.44 | 1 |
| Θ_OG_ animals | 1.47 | 1.37 | 1.44 | 1.51 | 1.57 | 5000.00 | 1 |
| λ animals | 0.04 | 0.03 | 0.04 | 0.05 | 0.08 | 5197.16 | 1 |
| Τ_OG_ animals | 0.02 | 0.01 | 0.02 | 0.03 | 0.04 | 5000.00 | 1 |
| Τ_REC_ animals | 0.08 | 0.05 | 0.07 | 0.1 | 0.14 | 5449.55 | 1 |
| Θ_0_ plants | 1.04 | 0.84 | 0.97 | 1.12 | 1.3 | 5000.00 | 1 |
| Θ_OG_ plants | 1.01 | 0.993 | 0.98 | 1.03 | 1.09 | 5103.35 | 1 |
| λ plants | 1.0 | 0.15 | 0.53 | 1.94 | 6.91 | 4831.69 | 1 |
| Τ_OG_ plants | 0.11 | 0.06 | 0.09 | 0.14 | 0.24 | 5000.00 | 1 |
| Τ_REC_ plants | 0.05 | 0.03 | 0.05 | 0.06 | 0.09 | 4874.01 | 1 |

Table S2. Functional traits of plant species observed in seed-dispersal interactions. Traits of plant species observed interacting in early recovery (ER), late recovery (LR) and old-growth (OG) forests. Crop mass is given in grams, fruit width in millimeters and height in meters. Plant numbers (#) correspond to those shown in Fig. 2C of the main text. Species are ordered from the lowest to the highest originality value.

| **#** | **Family** | **Species** | **CropMass** | **FruitWidth** | **Height** | **ER** | **LR** | **OG** |
| --- | --- | --- | --- | --- | --- | --- | --- | --- |
| 1 | Sapinadaceae | *Allophylus angustatus* | 244.80 | 8.66 | 19.00 |  | X |  |
| 2 | Myrtaceae | *Myrcia aliena* | 1334.25 | 13.83 | 15.00 |  |  | X |
| 3 | Arecaceae | *Prestoea ensiformis* | 788.77 | 11.90 | 13.25 |  |  | X |
| 4 | Meliaceae | *Guarea guidonia* | 560.00 | 14.24 | 42.00 |  |  | X |
| 5 | Melastomataceae | *Conostegia superba* | 567.31 | 8.98 | 8.62 | X | X | X |
| 6 | Moraceae | *Ficus pertusa* | 146.16 | 10.59 | 20.00 |  |  | X |
| 7 | Moraceae | *Brosimum guianense* | 450.00 | 15.77 | 40.00 |  |  | X |
| 8 | Moraceae | *Ficus brevibracteata* | 157.19 | 10.09 | 19.83 | X | X | X |
| 9 | Malvaceae | *Matisia longipes* | 1152.80 | 19.28 | 23.60 |  |  | X |
| 10 | Urticaceae | *Cecropia garciae* | NA | NA | 22.15 | X | X |  |
| 11 | Urticaceae | *Cecropia insignis* | 226.86 | 11.10 | 25.25 | X | X | X |
| 12 | Rubiaceae | *Palicourea timbiquensis* | 112.73 | 6.51 | 3.63 | X |  |  |
| 13 | Rubiaceae | *Isertia pittieri* | 1065.20 | 13.18 | 7.92 | X | X |  |
| 14 |  | *A-CR1303* | 36.90 | 5.54 | 6.25 | X |  |  |
| 15 | Fabaceae | *Hymenolobium heterocarpum* | NA | NA | 38.50 |  | X | X |
| 16 | Meliaceae | *Trichilia surinamensis* | 851.40 | 10.68 | 18.50 |  | X | X |
| 17 | Melastomataceae | *Miconia multiplicata* | 111.63 | 5.32 | 14.00 | X | X |  |
| 18 |  | *A-CR0601* | 377.60 | 11.53 | 31.00 |  | X |  |
| 19 | Myrtaceae | *Eugenia multirimosa* | 487.50 | 10.54 | 29.50 |  | X |  |
| 20 | Chrysobalanaceae | *Couepia subcordata* | 93.00 | 6.56 | 34.00 |  |  | X |
| 21 | Myristicaceae | *Virola reidii* | 2606.00 | 21.25 | 28.75 |  |  | X |
| 22 | Rubiaceae | *Pentagonia macrophylla* | 41.16 | 6.40 | 10.14 |  | X |  |
| 23 | Urticaceae | *Cecropia hispidissima* | 127.01 | 5.63 | 18.37 | X | X | X |
| 24 | Moraceae | *Brosimum utile* | 2084.68 | 22.11 | 33.06 |  |  | X |
| 25 | Melastomataceae | *Miconia centronioides* | 135.95 | 6.04 | 3.42 | X | X |  |
| 26 | Lauraceae | *Nectandra purpurea* | 856.70 | 10.88 | 25.00 |  | X |  |
| 27 | Urticaceae | *Pourouma guianensis* | 2507.78 | 18.12 | 34.17 |  |  | X |
| 28 | Meliaceae | *Guarea bullata* | 304.36 | 6.90 | 18.67 |  | X | X |
| 29 | Arecaceae | *Wettinia quinaria* | 3812.96 | 18.12 | 18.18 | X | X | X |
| 30 |  | *A-CR1803* | 988.00 | 7.17 | 4.00 | X |  |  |
| 31 | Melastomataceae | *Miconia oraria* | 157.36 | 5.40 | 6.61 | X | X | X |
| 32 | Myristicaceae | *Virola pavonis* | 2170.38 | 20.14 | 23.78 |  | X |  |
| 33 | Malvaceae | *Matisia bracteolosa* | 1600.00 | 15.58 | 46.00 |  |  | X |
| 34 | Rubiaceae | *Palicourea pseudaxillaris* | 215.74 | 5.19 | 6.56 | X | X |  |
| 35 | Staphyleaceae | *Turpinia occidentalis* | 864.68 | 9.44 | 24.60 |  | X |  |
| 36 | Arecaceae | *Bactris setulosa* | 1137.89 | 19.35 | 14.90 |  | X | X |
| 37 | Rubiaceae | *Palicourea paratinctoria* | 80.62 | 5.16 | 6.08 | X | X | X |
| 38 | Myristicaceae | *Otoba gracilipes* | 2756.52 | 15.16 | 35.00 |  |  | X |
| 39 | Chrysobalanaceae | *Licania glauca* | 1621.80 | 12.25 | 25.00 |  | X |  |
| 40 | Rutaceae | *Hortia brasiliana* | 1197.00 | 24.99 | 41.00 |  |  | X |
| 41 | Piperaceae | *Piperacea sp2* | 132.18 | 9.16 | 2.38 | X | X |  |
| 42 | Lecythidaceae | *Eschweilera decolorans* | 1976.25 | 25.97 | 56.00 |  |  | X |
| 43 | Hypericaceae | *Vismia baccifera* | 1704.24 | 9.48 | 13.00 | X | X |  |
| 44 | Cannabaceae | *Trema micrantha* | 309.30 | 16.23 | 17.45 | X | X |  |
| 45 | Burseraceae | *Protium ecuadorense* | 9859.85 | 27.66 | 32.63 |  |  | X |
| 46 | Salicaceae | *Banara guianensis* | 1426.79 | 9.11 | 14.76 | X | X |  |
| 47 | Myristicaceae | *Otoba novogranatensis* | 6567.92 | 25.40 | 30.07 | X | X | X |
| 48 | Verbanaceae | *Lantana sp.* | 86.59 | 3.45 | 4.30 | X |  |  |
| 49 |  | *A-PR3201* | 34.44 | 5.17 | 4.89 |  | X | X |
| 50 | Lauraceae | *Nectandra guadaripo* | 696.00 | 8.37 | 48.00 |  |  | X |
| 51 | Melastomataceae | *Miconia multiplicata* | 136.41 | 4.98 | 7.00 | X |  | X |
| 52 | Melastomataceae | *Miconia conocuatrecasii* | 22.60 | 4.10 | 6.20 | X | X | X |
| 53 | Bignoniaceae | *Tabebuia rosea* | 173.60 | 4.04 | 18.00 |  | X |  |
| 54 | Annonaceae | *Guatteria pittieri* | 258.30 | 4.61 | 24.00 | X | X |  |
| 55 | Arecaceae | *Iriartea deltoidea* | 5035.97 | 11.90 | 24.88 |  |  | X |
| 56 |  | *A-CR1302* | 27.63 | 6.67 | 2.00 | X |  |  |
| 57 | Euphorbiaceae | *Conceveiba martiana* | 2677.50 | 33.09 | 30.25 |  | X | X |
| 58 | Euphorbiaceae | *Tetrorchidium andinum* | 598.00 | 6.29 | 28.00 |  |  | X |
| 59 |  | *A-OG49002* | 30.02 | 3.64 | 5.50 |  | X | X |
| 60 | Malvaceae | *Apeiba membranacea* | 12605.36 | 47.54 | 44.50 |  |  | X |
| 61 |  | *A-CR0501* | 107.03 | 13.39 | 2.00 | X |  |  |
| 62 |  | *A-PA5601* | 32.39 | 4.80 | 0.75 | X |  |  |
| 63 |  | *A-CR0101* | 131.25 | 15.48 | 2.00 | X |  |  |
| 64 | Sabiaceae | *Meliosma herbertii* | 279.58 | 24.69 | 17.00 |  | X |  |
| 65 |  | *A-PR2101* | 924.12 | 21.90 | 1.00 |  | X | X |
| 66 | Myristicaceae | *Osteophloeum platyspermum* | 51999.23 | 30.25 | 47.00 |  |  | X |
| 67 | Melastomataceae | *Miconia montana* | 9.63 | 1.63 | 12.89 |  | X |  |
| 68 | Burseraceae | *Protium colombianum* | 520.60 | 38.85 | 52.00 |  |  | X |
| 69 | Malvaceae | *Ficus maxima* | 13626.00 | 33.69 | 46.00 |  | X |  |
| 70 |  | *A-CR1601* | 49.00 | 2.20 | 1.25 | X |  |  |
| 71 | Arecaceae | *Socratea exorrhiza* | 131.33 | 28.19 | 26.50 |  |  | X |
| 72 | Melastomataceae | *Miconia sp3* | 16.01 | 4.74 | 2.18 |  | X | X |
| 73 | Caryocaraceae | *Caryocar glabrum* | 13892.57 | 12.78 | 43.00 |  |  | X |
| 74 | Phyllanthaceae | *Hieronyma alchorneoides* | 333.88 | 3.31 | 23.13 | X | X | X |
| 75 | Melastomataceae | *Miconia sp5* | 4.92 | 4.43 | 1.17 | X |  |  |
| 76 | Rubiaceae | *Coussarea latifolia* | 204.75 | 31.20 | 13.00 | X |  | X |
| 77 |  | *A-CR1111* | 19.74 | 5.39 | 1.00 |  | X | X |
| 78 | Lecythidaceae | *Grias peruviana* | 5967.00 | 58.87 | 6.50 | X |  |  |
| 79 | Meliaceae | *Carapa guianensis* | 4543.70 | 94.17 | 23.50 |  |  | X |
| 80 |  | *A-PR2601* | 11.40 | 8.12 | 1.00 |  | X |  |
| 81 |  | *A-OG4701* | 931.71 | 59.95 | 3.00 |  |  | X |

Table S3. Functional traits of animal species observed in seed-dispersal interactions. Traits of animal species observed interacting in early recovery (ER), late recovery (LR) and old-growth (OG) forests. Body mass is given in grams and gape width in millimeters. HWI stands for hand-wing index. Animal numbers (#) correspond to those shown in Fig. 2B of the main text. Species are ordered from the lowest to the highest originality value.

| **#** | **Order** | **Species** | **BodyMass** | **GapeWidth** | **HWI** | **ER** | **LR** | **OG** |
| --- | --- | --- | --- | --- | --- | --- | --- | --- |
| 1 | Galliformes | *Penelope ortoni* | 872.00 | 8.68 | 12.20 |  |  | X |
| 2 | Passeriformes | *Tityra inquisitor* | 43.10 | 11.63 | 26.48 |  | X |  |
| 3 | Passeriformes | *Lipaugus unirufus* | 82.10 | 7.18 | 20.45 | X | X | X |
| 4 | Passeriformes | *Sicalis flaveola* | 16.90 | 5.40 | 20.61 | X |  |  |
| 5 | Passeriformes | *Thraupis episcopus* | 35.00 | 6.40 | 22.85 | X |  |  |
| 6 | Passeriformes | *Thraupis palmarum* | 39.00 | 5.87 | 21.85 | X | X |  |
| 7 | Passeriformes | *Heterospingus xanthopygius* | 38.80 | 6.38 | 20.27 | X | X | X |
| 8 | Piciformes | *Capito squamatus* | 59.30 | 8.30 | 19.35 | X | X | X |
| 9 | Passeriformes | *Myiozetetes cayanensis* | 25.90 | 5.35 | 15.43 | X |  |  |
| 10 | Passeriformes | *Turdus maculirostris* | 69.60 | 4.87 | 18.05 |  | X |  |
| 11 | Passeriformes | *Mitrospingus cassinii* | 40.40 | 5.38 | 12.55 | X |  |  |
| 12 | Passeriformes | *Tangara cyanicollis* | 17.00 | 4.46 | 21.93 | X |  |  |
| 13 | Passeriformes | *Euphonia laniirostris* | 15.00 | 5.58 | 23.39 | X | X |  |
| 14 | Rodentia | *Heteromys australis* | 267.50 | 12.17 | NA | X | X | X |
| 15 | Passeriformes | *Cacicus cela* | 85.50 | 7.94 | 27.22 |  |  | X |
| 16 | Passeriformes | *Tangara gyrola* | 21.00 | 4.83 | 21.08 | X | X |  |
| 17 | Passeriformes | *Saltator maximus* | 47.60 | 8.86 | 16.94 | X |  |  |
| 18 | Passeriformes | *Tangara palmeri* | 32.30 | 5.20 | 21.28 | X | X | X |
| 19 | Passeriformes | *Tangara larvata* | 20.00 | 4.60 | 21.45 | X | X |  |
| 20 | Passeriformes | *Dacnis berlepschi* | 12.80 | 3.63 | 19.08 | X |  |  |
| 21 | Passeriformes | *Saltator coerulescens* | 54.90 | 8.83 | 14.51 | X |  |  |
| 22 | Passeriformes | *Sporophila corvina* | 10.60 | 5.28 | 17.80 | X |  |  |
| 23 | Passeriformes | *Saltator grossus* | 44.20 | 10.30 | 18.57 | X | X |  |
| 24 | Passeriformes | *Coereba flaveola* | 10.00 | 3.60 | 19.20 | X |  |  |
| 25 | Passeriformes | *Myiobius barbatus* | 11.00 | 4.26 | 16.82 | X |  |  |
| 26 | Coraciiformes | *Baryphthengus martii* | 165.00 | 11.65 | 16.11 | X |  |  |
| 27 | Passeriformes | *Tangara johannae* | 20.00 | 4.80 | 24.25 | X | X | X |
| 28 | Cuculiformes | *Piaya cayana* | 102.00 | 8.79 | 10.98 | X |  |  |
| 29 | Passeriformes | *Ramphocelus flammigerus* | 33.00 | 8.22 | 15.08 | X | X |  |
| 30 | Passeriformes | *Querula purpurata* | 107.40 | 12.48 | 17.60 | X | X | X |
| 31 | Passeriformes | *Tangara lavinia* | 24.00 | 4.25 | 23.90 | X | X | X |
| 32 | Passeriformes | *Euphonia xanthogaster* | 13.00 | 4.83 | 22.38 | X | X | X |
| 33 | Passeriformes | *Manacus manacus* | 16.70 | 4.42 | 15.91 | X | X |  |
| 34 | Passeriformes | *Tangara florida* | 19.30 | 4.30 | 24.50 | X | X | X |
| 35 | Passeriformes | *Chlorophanes spiza* | 19.00 | 4.46 | 24.74 | X | X | X |
| 36 | Galliformes | *Penelope purpurascens* | 2060.00 | 10.14 | 12.03 |  | X |  |
| 37 | Passeriformes | *Arremon aurantiirostris* | 34.50 | 6.57 | 11.31 | X |  |  |
| 38 | Tinamiformes | *Tinamus major* | 506.60 | 5.15 | 18.67 |  |  | X |
| 39 | Passeriformes | *Dacnis cayana* | 13.00 | 3.70 | 24.70 | X | X |  |
| 40 | Passeriformes | *Euphonia saturata* | 13.30 | 3.94 | 22.23 |  | X |  |
| 41 | Psittaciformes | *Amazona farinosa* | 626.00 | 20.54 | 27.86 |  |  | X |
| 42 | Passeriformes | *Dacnis venusta* | 16.10 | 3.81 | 26.30 | X |  |  |
| 43 | Passeriformes | *Machaeropterus deliciosus* | 12.70 | 3.84 | 20.53 |  | X |  |
| 44 | Psittaciformes | *Amazona autumnalis* | 416.00 | 16.43 | 31.04 |  |  | X |
| 45 | Passeriformes | *Chlorothraupis olivacea* | 39.00 | 8.23 | 18.30 |  |  | X |
| 46 | Galliformes | *Odontophorus erythrops* | 335.00 | 8.86 | 27.05 |  | X | X |
| 47 | Trogoniformes | *Trogon comptus* | 114.00 | 9.85 | 33.43 |  | X | X |
| 48 | Passeriformes | *Tachyphonus delatrii* | 18.00 | 4.92 | 15.60 | X | X | X |
| 49 | Passeriformes | *Ceratopipra mentalis* | 15.00 | 4.38 | 16.49 | X | X | X |
| 50 | Passeriformes | *Sapayoa aenigma* | 20.80 | 7.70 | 19.03 | X |  | X |
| 51 | Passeriformes | *Arremonops conirostris* | 35.90 | 6.22 | 8.15 | X |  |  |
| 52 | Passeriformes | *Tachyphonus luctuosus* | 13.00 | 4.88 | 17.64 |  | X |  |
| 53 | Coraciiformes | *Electron platyrhynchum* | 73.00 | 14.80 | 17.70 |  | X |  |
| 54 | Rodentia | *Hoplomys gymnurus* | 240.00 | 23.08 | NA |  | X | X |
| 55 | Passeriformes | *Cyanocompsa cyanoides* | 32.50 | 10.21 | 13.77 | X |  |  |
| 56 | Passeriformes | *Rhytipterna holerythra* | 36.80 | 7.08 | 16.50 |  |  | X |
| 57 | Trogoniformes | *Trogon chionurus* | 89.70 | 10.92 | 35.28 |  | X |  |
| 58 | Passeriformes | *Cyanerpes caeruleus* | 12.00 | 3.44 | 25.49 | X |  | X |
| 59 | Passeriformes | *Ornithion brunneicapillus* | 7.10 | 3.03 | 12.65 | X |  |  |
| 60 | Passeriformes | *Cephalopterus penduliger* | 338.00 | 18.42 | 14.38 |  | X | X |
| 61 | Passeriformes | *Catharus ustulatus* | 30.30 | 4.41 | 30.61 | X | X |  |
| 62 | Psittaciformes | *Pyrilia pulchra* | 150.00 | 10.80 | 36.50 |  |  | X |
| 63 | Passeriformes | *Lepidothrix coronata* | 8.30 | 3.66 | 17.88 | X | X | X |
| 64 | Passeriformes | *Chrysothlypis salmoni* | 12.50 | 3.90 | 17.48 |  | X | X |
| 65 | Rodentia | *Proechimys semispinosus* | 360.50 | 26.80 | NA | X | X | X |
| 66 | Psittaciformes | *Pyrrhura melanura* | 70.50 | 10.72 | 42.05 |  |  | X |
| 67 | Piciformes | *Pteroglossus torquatus* | 219.10 | 22.88 | 15.69 | X | X | X |
| 68 | Tinamiformes | *Crypturellus berlepschi* | 516.80 | 5.34 | 23.30 |  | X |  |
| 69 | Didelphimorphia | *Didelphis marsupialis* | 1091.16 | 31.82 | NA |  | X |  |
| 70 | Columbiformes | *Leptotila verreauxi* | 146.90 | 3.63 | 22.15 | X |  |  |
| 71 | Tinamiformes | *Tinamus major* | 1026.20 | 5.20 | 20.94 |  | X | X |
| 72 | Galliformes | *Rhynchortyx cinctus* | 150.00 | 7.94 | 36.25 |  |  | X |
| 73 | Rodentia | *Notosciurus granatensis* | 250.00 | 33.40 | NA | X | X | X |
| 74 | Rodentia | *Dasyprocta punctata* | 2674.98 | 37.22 | NA |  | X | X |
| 75 | Primates | *Cebus capucinus* | 2984.44 | NA | NA | X | X | X |
| 76 | Columbiformes | *Geotrygon montana* | 133.90 | 3.48 | 27.10 |  | X | X |
| 77 | Columbiformes | *Geotrygon purpurata* | 129.00 | 3.23 | 26.28 |  | X | X |
| 78 | Passeriformes | *Masius chrysopterus* | 11.50 | 3.82 | 12.80 |  |  | X |
| 79 | Rodentia | *Cuniculus paca* | 8172.55 | NA | NA |  |  | X |
| 80 | Piciformes | *Ramphastos brevis* | 412.00 | 32.88 | 14.35 | X | X | X |
| 81 | Psittaciformes | *Ara ambiguus* | 1300.00 | 32.34 | 39.18 |  |  | X |
| 82 | Primates | *Alouatta palliata* | 6120.00 | 60.43 | NA |  |  | X |
| 83 | Primates | *Ateles fusciceps* | 9050.00 | 55.93 | NA |  |  | X |
| 84 | Piciformes | *Ramphastos ambiguus* | 651.70 | 38.25 | 10.13 |  | X | X |
| 85 | Columbiformes | *Patagioenas goodsoni* | 176.00 | 3.98 | 34.20 |  | X | X |
| 86 | Columbiformes | *Patagioenas subvinacea* | 162.50 | 3.75 | 34.33 |  | X | X |
| 87 | Columbiformes | *Leptotrygon veraguensis* | 155.00 | 2.64 | 27.15 | X |  |  |
| 88 | Artiodactyla | *Tayassu pecari* | 32233.69 | 91.96 | NA |  | X | X |

Table S4. Seed-dispersal interactions across habitat types. List of observed interactions in each habitat type: early recovery (ER), late recovery (LR), and old-growth (OG) forests. Interaction numbers (#) correspond to those shown in Fig. 2A of the main text. Interactions are ordered from the lowest to the highest originality value. Only interactions observed at least six times were included in Fig. S2A (left) and in this table.

| **#** | **Interaction** | **ER** | **LR** | **OG** |
| --- | --- | --- | --- | --- |
| 1 | *Cecropia hispidissima.Myiobius barbatus* | X |  |  |
| 2 | *A-CR1803.Euphonia xanthogaster* | X |  |  |
| 3 | *Conostegia superba.Manacus manacus* | X |  |  |
| 4 | *Miconia oraria.Myiozetetes cayanensis* | X |  |  |
| 5 | *Cecropia insignis.Tangara gyrola* |  | X |  |
| 6 | *Cecropia insignis.Heterospingus xanthopygius* |  | X | X |
| 7 | *Miconia oraria.Thraupis episcopus* | X |  |  |
| 8 | *Guarea guidonia.Tachyphonus delatrii* |  |  | X |
| 9 | *A-CR1303.Tangara gyrola* | X |  |  |
| 10 | *Prestoea ensiformis.Lipaugus unirufus* |  |  | X |
| 11 | *Palicourea timbiquensis.Manacus manacus* | X |  |  |
| 12 | *Miconia oraria.Tangara cyanicollis* | X |  |  |
| 13 | *A-CR1303.Tachyphonus delatrii* | X |  |  |
| 14 | *A-CR1303.Tangara cyanicollis* | X |  |  |
| 15 | *Tabebuia rosea.Euphonia xanthogaster* |  | X |  |
| 16 | *Palicourea timbiquensis.Tachyphonus delatrii* | X |  |  |
| 17 | *A-CR1303.Mitrospingus cassinii* | X |  |  |
| 18 | *Hieronyma alchorneoides.Ateles fusciceps* |  |  | X |
| 19 | *Cecropia hispidissima.Tangara gyrola* | X |  |  |
| 20 | *Conostegia superba.Tachyphonus delatrii* |  | X |  |
| 21 | *Miconia oraria.Tangara gyrola* | X | X |  |
| 22 | *Cecropia insignis.Tachyphonus delatrii* |  | X | X |
| 23 | *Cecropia hispidissima.Tangara palmeri* | X | X |  |
| 24 | *Cecropia hispidissima.Thraupis episcopus* | X |  |  |
| 25 | *Lantana sp..Euphonia xanthogaster* | X |  |  |
| 26 | *Miconia oraria.Manacus manacus* | X | X |  |
| 27 | *Guatteria pittieri.Saltator grossus* | X | X |  |
| 28 | *Cecropia hispidissima.Thraupis palmarum* | X |  |  |
| 29 | *Cecropia insignis.Tangara johannae* |  | X | X |
| 30 | *Miconia oraria.Saltator coerulescens* | X |  |  |
| 31 | *Miconia oraria.Querula purpurata* | X | X |  |
| 32 | *Miconia oraria.Tachyphonus delatrii* | X | X |  |
| 33 | *Cecropia hispidissima.Ramphocelus flammigerus* | X |  |  |
| 34 | *Cecropia hispidissima.Tachyphonus delatrii* | X |  | X |
| 35 | *Lantana sp..Dacnis berlepschi* | X |  |  |
| 36 | *Lantana sp..Mitrospingus cassinii* | X |  |  |
| 37 | *Ficus pertusa.Rhytipterna holerythra* |  |  | X |
| 38 | *A-CR1303.Ramphocelus flammigerus* | X |  |  |
| 39 | *Cecropia insignis.Tangara florida* |  | X | X |
| 40 | *Miconia centronioides.Capito squamatus* |  | X |  |
| 41 | *Cecropia insignis.Tangara lavinia* |  | X | X |
| 42 | *Miconia centronioides.Tangara palmeri* |  | X |  |
| 43 | *Miconia centronioides.Tachyphonus delatrii* | X | X |  |
| 44 | *Ficus pertusa.Tachyphonus delatrii* |  |  | X |
| 45 | *Miconia oraria.Ceratopipra mentalis* | X | X | X |
| 46 | *Lantana sp..Ramphocelus flammigerus* | X |  |  |
| 47 | *Miconia oraria.Ramphocelus flammigerus* | X | X |  |
| 48 | *Couepia subcordata.Tachyphonus delatrii* |  |  | X |
| 49 | *Miconia oraria.Lepidothrix coronata* | X | X | X |
| 50 | *Vismia baccifera.Thraupis episcopus* | X |  |  |
| 51 | *Lantana sp..Tangara lavinia* | X |  |  |
| 52 | *Pourouma guianensis.Cacicus cela* |  |  | X |
| 53 | *Brosimum utile.Cacicus cela* |  |  | X |
| 54 | *Miconia conocuatrecasii.Euphonia xanthogaster* |  | X |  |
| 55 | *Lantana sp..Cyanerpes caeruleus* | X |  |  |
| 56 | *Miconia conocuatrecasii.Tangara lavinia* |  | X |  |
| 57 | *Turpinia occidentalis.Cebus capucinus* |  | X |  |
| 58 | *Brosimum utile.Proechimys semispinosus* |  |  | X |
| 59 | *Brosimum utile.Dasyprocta punctata* |  |  | X |
| 60 | *Conostegia superba.Lepidothrix coronata* |  |  | X |
| 61 | *Cecropia hispidissima.Dacnis venusta* | X |  |  |
| 62 | *Wettinia quinaria.Hoplomys gymnurus* |  | X | X |
| 63 | *Nectandra purpurea.Cephalopterus penduliger* |  | X |  |
| 64 | *Pourouma guianensis.Ateles fusciceps* |  |  | X |
| 65 | *Miconia oraria.Catharus ustulatus* | X | X |  |
| 66 | *Cecropia insignis.Pteroglossus torquatus* |  |  | X |
| 67 | *A-CR1601.Tachyphonus delatrii* | X |  |  |
| 68 | *Miconia montana.Tachyphonus delatrii* |  | X |  |
| 69 | *Eugenia multirimosa.Trogon comptus* |  | X |  |
| 70 | *Miconia multiplicata.Tachyphonus delatrii* |  |  | X |
| 71 | *Conostegia superba.Ramphastos brevis* |  | X |  |
| 72 | *Miconia conocuatrecasii.Tachyphonus delatrii* |  | X | X |
| 73 | *Pourouma guianensis.Pyrilia pulchra* |  |  | X |
| 74 | *Nectandra purpurea.Ramphastos brevis* |  | X |  |
| 75 | *Miconia multiplicata.Geotrygon purpurata* |  | X |  |
| 76 | *Guatteria pittieri.Ramphastos ambiguus* |  | X |  |
| 77 | *A-OG49002.Tachyphonus delatrii* |  |  | X |
| 78 | *Virola reidii.Ramphastos ambiguus* |  |  | X |
| 79 | *Conceveiba martiana.Ara ambiguus* |  |  | X |
| 80 | *Conostegia superba.Ramphastos ambiguus* |  | X |  |
| 81 | *Otoba novogranatensis.Pyrrhura melanura* |  |  | X |
| 82 | *Turpinia occidentalis.Ramphastos ambiguus* |  | X |  |
| 83 | *Cecropia insignis.Ramphastos ambiguus* |  | X |  |
| 84 | *Nectandra purpurea.Ramphastos ambiguus* |  | X |  |
| 85 | *Miconia conocuatrecasii.Patagioenas goodsoni* |  | X |  |
| 86 | *Banara guianensis.Patagioenas goodsoni* |  | X |  |
